## Supplementary material for "Differential Conditioning Produces Merged Long-term Memory in *Drosophila*": Figure Supplement

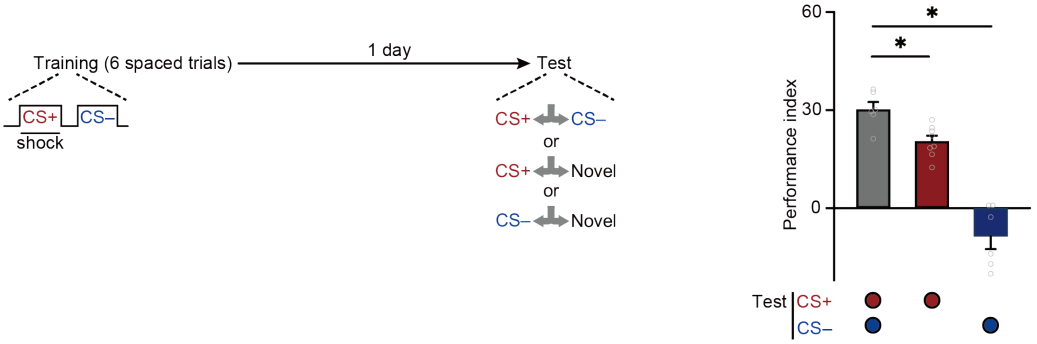


**Figure 1-figure supplement 1. Repetitive Spaced Training Forms Complementary LTMs.** Multi-trial spaced training (six trials of training with 15-min interval) induced 24-h aversive memory to CS+ and approach memory to CS– (n = 6-8). All data shown are presented as mean ± SEM. *$p < 0.05$.

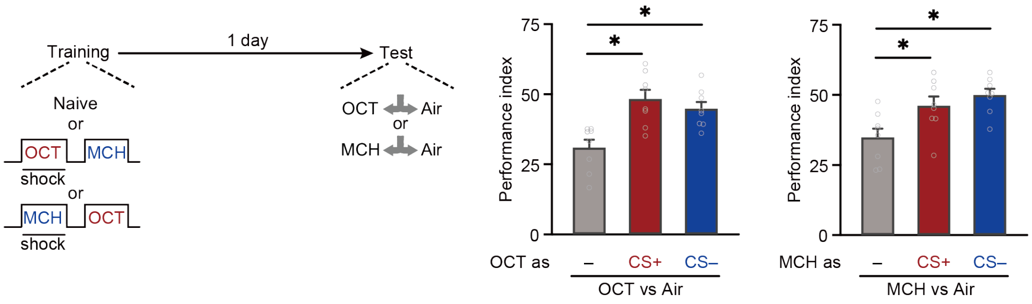


**Figure 1-figure supplement 2. Single-trial Differential Conditioning Increases CS+ and CS– Odor Avoidances.** Odor avoidances of OCT and MCH were significantly increased when they were used as CS+ or CS– during training (n = 8). All data shown are presented as mean ± SEM. *$p < 0.05$.

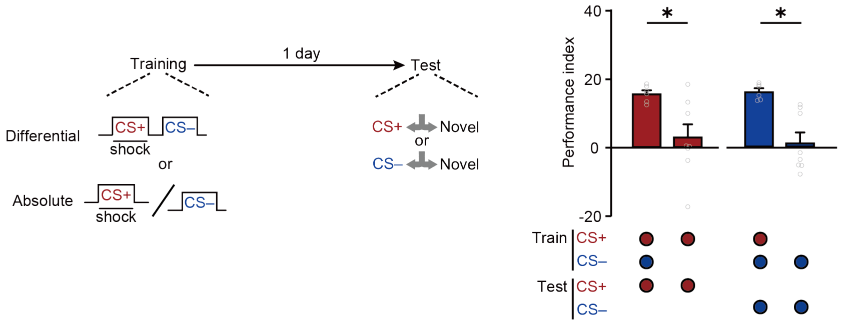


**Figure 2-figure supplement 1. Absolute Trainings Fail to Induce 24-h Avoidances.** Training without CS+ or CS– abolished the 24-h avoidance to CS– or CS+ (n = 6-8). All data shown are presented as mean ± SEM. *$p < 0.05$.

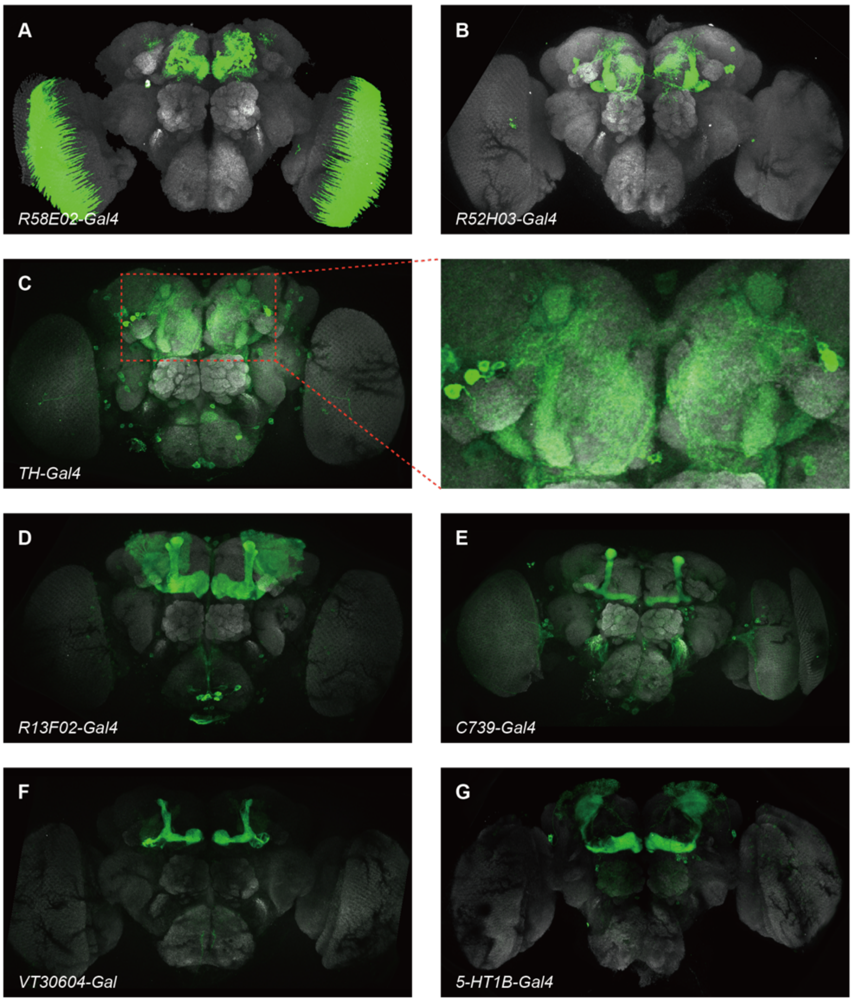


**Figure 3-figure supplement 1. The Expression Patterns of GAL4 Lines Used in Figure 3.** Panels show GFP expression driven by the relevant GAL4 (green) and general neuropil stained with an antibody to the presynaptic marker nc82 (grey). **(A)** *R58E02-Gal4* labels all PAM DANs. **(B)** *R52H03-Gal4* labels all PPL1 DANs. **(C)** *TH-Gal4* broadly labels all PPL1 DANs. **(D)** *R13F02-Gal4* labels all KCs in MB. **(E)** *C739-Gal4* labels αβ KCs. **(F)** *VT20604-Gal4* labels α′β′ KCs. **(G)** *5-HT1B-Gal4* labels γ KCs.

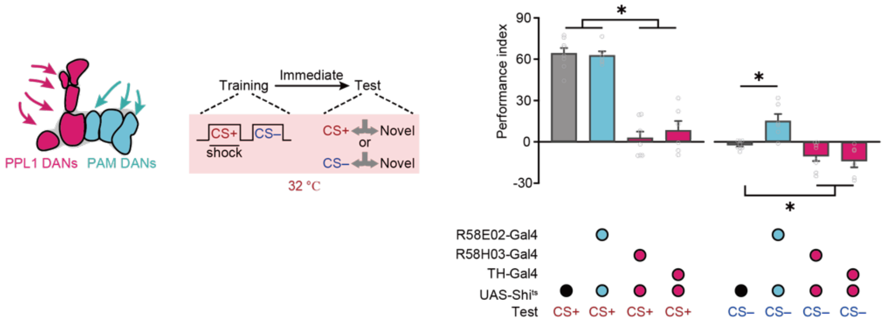


**Figure 3-figure supplement 2. PPL1 and PAM DANs Play Different Roles in CS+ and CS– Memory Encoding.** Blocking PAM DANs did not affect the 3-min avoidance to CS+ but induced a significant avoidance to CS–, whereas blocking PPL1 DANs abolished the 3-min avoidance to CS+ but increases the 3-min approach to CS– (n = 6-8). All data shown are presented as mean ± SEM. *$p < 0.05$.

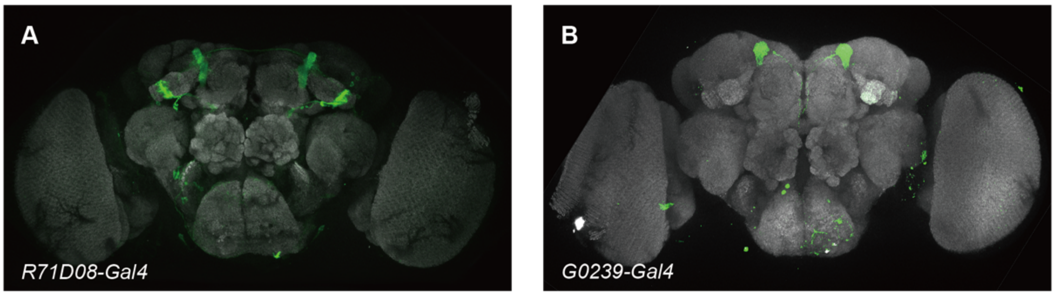


**Figure 4-figure supplement 1. The Expression Patterns of GAL4 Lines Used in Figure 4.** Panels show GFP expression driven by the relevant GAL4 (green) and general neuropil stained with an antibody to the presynaptic marker nc82 (grey). **(A)** *R71E08-Gal4* labels α2sc MBONs. **(B)** *G0239-Gal4* labels α3 MBONs.

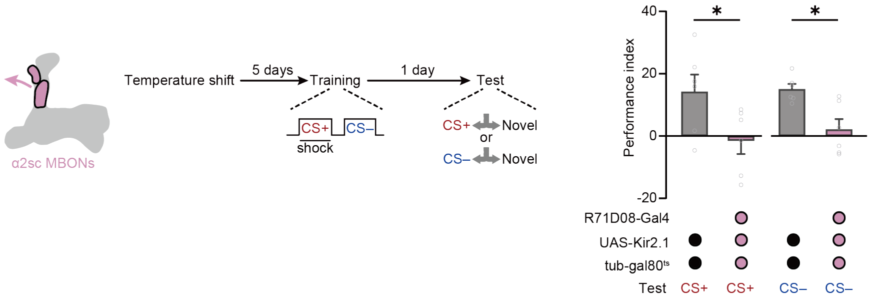


**Figure 4-figure supplement 2. Retrieving the mLTM Relies on the Neural Activity of α2sc MBONs.** Blocking the neural activity of α2sc MBONs using *R71D08-Gal4>UAS-Kir2.1;tub-gal80*$^{ts}$ abolished the mLTM expression (n = 6). Flies were moved to high temperature (32°C) 5 days before training for the induction of Kir2.1. All data shown are presented as mean ± SEM. *p < 0.05.

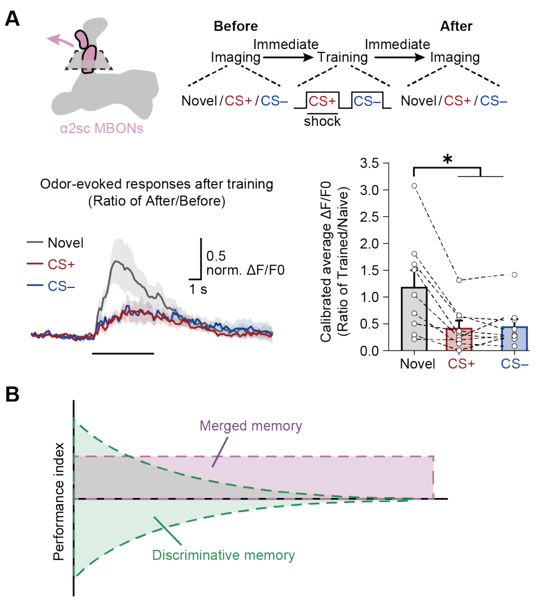


**Figure 5-figure supplement 1. The Depression of Odor-evoked Responses in α2sc MBONs Can Be Observed Immediately after Training. (A)** Top: training and imaging protocols. The odor-evoked responses of novel odor, CS+, and CS– were sequentially recorded before and after training. Down: the depressed CS+ and CS– odor-evoked responses were recorded immediately after training (n = 9). All post-training responses were calibrated to the average responses of corresponding in same flies before training. **(B)** Model of memory components induced by single-trial differential conditioning. After training, two categories of memories are formed: the short-lasting discriminative memory guiding avoidance to CS+ and approach to CS–; and the long-lasting merged memory guiding avoidance to both CS+ and CS–. All data shown are presented as mean ± SEM. *$p < 0.05$.
